# Lamin B1 and LAP2β resist cytoskeletal force to maintain lamin A/C meshwork organization and preserve nuclear integrity

**DOI:** 10.1101/2025.02.16.636527

**Authors:** Yewande Alabi, Vasilisa Aksenova, Alexei Arnaoutov, Harold Marin, Mary Dasso, Abigail Buchwalter

## Abstract

The nuclear lamins are extremely long-lived proteins in most cell types. As a consequence, lamin function cannot be effectively dissected with temporal precision using standard knock-down approaches. Here, we apply the auxin-inducible degron (AID) system to rapidly deplete each lamin isoform within one cell cycle and reveal the immediate impacts of lamin loss on the nucleus. Surprisingly, neither acute lamin A/C (LA/C), lamin B1 (LB1), nor lamin B2 (LB2) depletion altered nuclear shape or induced nuclear blebbing, indicating that acute lamin loss is not sufficient to alter nuclear morphology. LB1 depletion is immediately followed by LA/C meshwork disorganization due to actin cytoskeletal forces on the lamina, yet neither LA/C nor LB1 depletion induced nuclear rupturing. We found that the abundant inner nuclear membrane protein LAP2β protects nuclear integrity in the absence of LB1, as depletion of both LB1 and LAP2β induced severe LA/C disorganization and frequent nuclear rupturing. Depolymerization of the actin cytoskeleton prevents nuclear rupture in LAP2β- and LB1-depleted nuclei. We conclude that both LB1 and LAP2β resist cytoskeletal force to maintain regular lamin A/C meshwork organization and preserve nuclear integrity.

## Introduction

The nuclear lamina (NL) is a filamentous protein meshwork at the nuclear periphery that underlies the inner nuclear membrane (INM). In mammals, two types of lamin proteins make up the NL: A-type lamins, consisting of lamin A and its splice isoform lamin C (LA/C, encoded by the *LMNA* gene), and B-type lamins, consisting of lamin B1 and lamin B2 (LB1 and LB2, encoded by *LMNB1* and *LMNB2* respectively). Insights from mouse and human genetics imply unique functions for each of the mammalian lamin isoforms. LA/C knockout (KO) mice die shortly after birth with severe cardiomyopathy (Kubben *et al*., 2011), and mutations to the human *LMNA* gene cause dilated cardiomyopathy, muscular dystrophy, lipodystrophy, progeria, and other syndromes (Schreiber and Kennedy, 2013). Both LB1 and LB2 KO mice die shortly after birth due to severe neurodevelopmental defects (Coffinier *et al*., 2010, 2011). However, neither LB1 nor LB2 can rescue the KO of the other (Lee *et al*., 2014), suggesting that B-type lamins have similar yet distinct roles.

Much of what we have learned about lamin function has come from KO or RNA-mediated knockdown (KD) approaches. However, the lamins are long-lived proteins, especially in non-dividing cells (Hasper *et al*., 2023). Therefore, any acute KO or KD approaches will have a long latency period, over multiple cell cycles, before an effect on lamin abundance will become apparent. As a result, it has not been possible to disrupt lamin function with temporal precision to reveal the immediate and direct impacts of lamin loss on nuclear function.

The lamins have been implicated in the regulation of gene expression and chromatin organization. Lamina-associated domains (LADs) of heterochromatin are enriched underneath the NL by interactions with the lamins and/or with lamin-associated proteins (Steensel and Belmont, 2017). LADs are transcriptionally repressive (Reddy *et al*., 2008; Leemans *et al*., 2019) and silence lineage-irrelevant genes during differentiation (Poleshko *et al*., 2017; Shah *et al*., 2021). In a role separate from LADs, a nucleoplasmic pool of LA/C can bind to active enhancers and promoters and is associated with active transcription (Gesson *et al*., 2016; Ikegami *et al*., 2020).

As the cell’s largest and most rigid organelle, the nucleus is constantly subjected to force (Miroshnikova and Wickström, 2021). The NL protects the nucleus from external force, and loss of lamin isoforms is associated with nuclear morphology defects. For instance, LA/C loss has been linked to decreased nuclear circularity and increased nuclear deformability(Sullivan *et al*., 1999a; Broers *et al*., 2004a; Lammerding *et al*., 2006), while LB1 loss has been linked to nuclear blebbing (Lammerding *et al*., 2006; Nmezi *et al*., 2019). The inability of the nucleus to withstand force leads to nuclear envelope (NE) rupture, which mixes nuclear and cytoplasmic contents and induces DNA damage (Maciejowski and Hatch, 2020). Both LA/C and LB1 have been implicated in preserving nuclear integrity (Vos *et al*., 2011; Vargas *et al*., 2012; Robijns *et al*., 2016; Chen *et al*., 2018; Earle *et al*., 2019). Lamin-associated INM proteins including LAP2β and LBR have also been linked to regulation of nuclear integrity (Maciejowski *et al*., 2015; Chen *et al*., 2021a; Baird *et al*., 2023).

Here, we leverage the auxin-inducible degron system(Nishimura *et al*., 2009; Morawska and Ulrich, 2013) to better understand the acute consequences of lamin loss of function. We demonstrate that we can rapidly deplete LA/C, LB1, and LB2 to undetectable levels within one cell cycle. Acute depletion of each lamin isoform has minor effects on gene expression and does not immediately induce changes to nuclear morphology. However, we show that LB1 depletion coincides with the opening of gaps in the LA/C meshwork and demonstrate that this effect depends on the actin cytoskeleton. Surprisingly, however, neither LA/C nor LB1-depleted cells undergo nuclear rupture. Instead, we determine that the highly expressed INM protein LAP2β and LB1 each organize the nuclear lamina and prevent nuclear rupture.

## Results

### Validation of auxin-inducible degrons for acute lamin depletion

To query the immediate impact of lamin depletion on nuclear function, we used CRISPR/Cas9 to biallelically insert a mNeonGreen (mNG) fluorescent protein and an auxin-inducible degron tag (micro-AID, (Morawska and Ulrich, 2013) immediately after the start codon of the *LMNA*, *LMNB1*, or *LMNB2* loci in the human colorectal adenocarcinoma cell line DLD1 (Figure 1B). To express the Tir1 ligase that mediates the ubiquitination of AID-tagged proteins in the presence of the plant hormone auxin, we used CRISPR/Cas9 to insert the sequence encoding OsTir1 into the 3’ end of the ubiquitously expressed *RCC1* gene, separated from the *RCC1* coding sequence by a cleavable P2A sequence (Aksenova *et al*., 2020). We confirmed the success of these edits by genotyping (Supplemental Figure 1A-C) and by Western blotting (Figure 1C-E), which indicates the expected molecular weight shifts corresponding to tag insertion. Tagging each lamin isoform moderately decreased both transcript and protein abundance (Supplemental Figure 3C; Figure 1C-E), implying transcriptional and post-transcriptional effects on the lamin proteins due to editing of their genomic loci and/or leaky degradation. Nevertheless, each tagged lamin isoform localizes normally to the nuclear periphery in the absence of auxin (Figure 2A). The addition of auxin to the cell culture medium reduced each lamin isoform to undetectable levels within 16 hours, or less than one cell cycle (Figure 1C-E). Auxin-mediated degradation is thus an effective way to rapidly deplete and probe the functions of these long-lived nuclear proteins.

**Figure 1.**
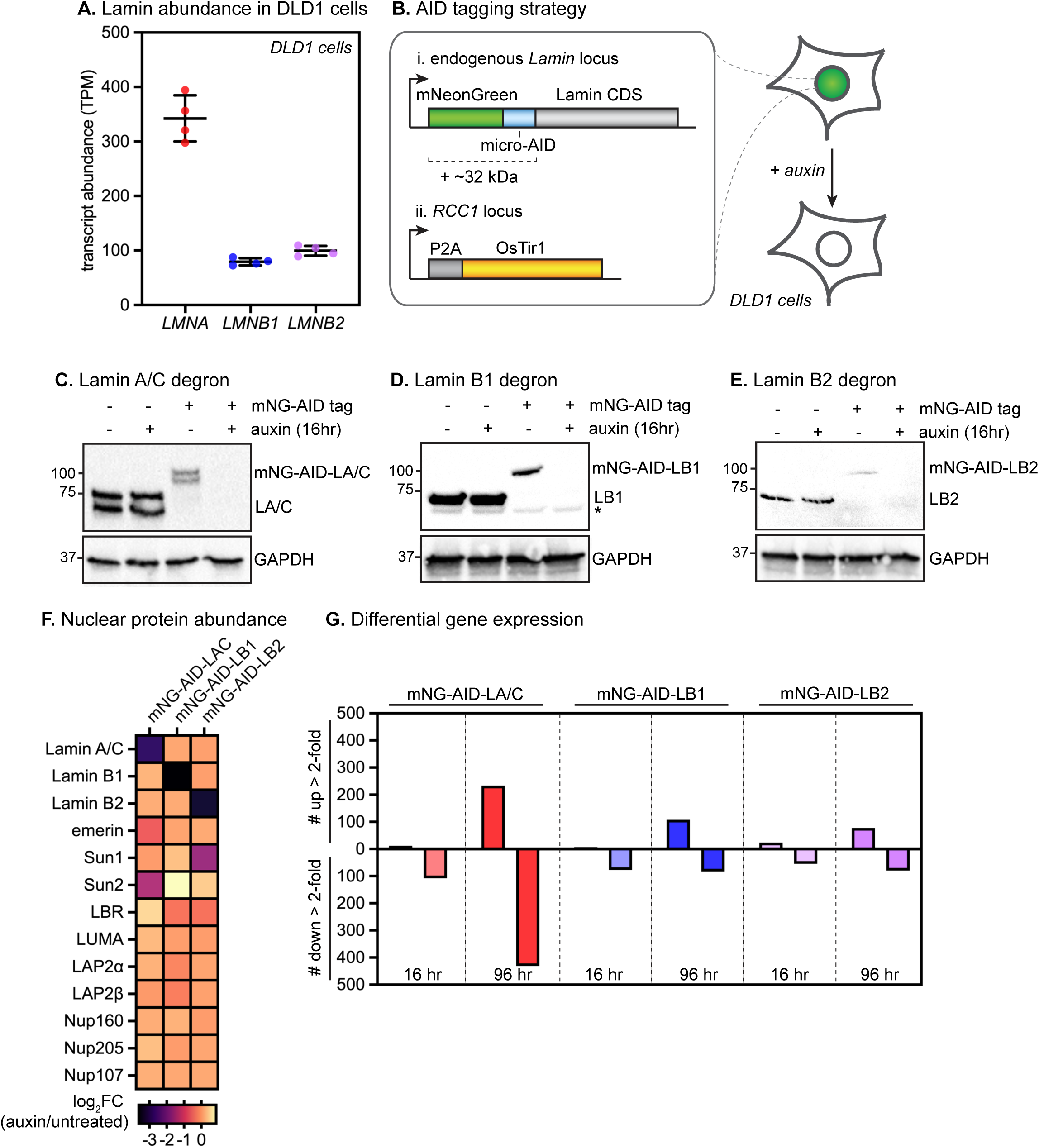
Auxin-inducible degradation of lamin A/C, lamin B1, and lamin B2 in DLD1 cells. (A) Relative expression levels of *LMNA, LMNB1,* and *LMNB2* genes in DLD1 cells. (B) Schematic of two-component auxin-inducible degron system, consisting of (i) N-terminal bi-allelic insertion of an mNeonGreen fluorescent protein and a micro-AID tag into the *LMNA, LMNB1,* or *LMNB2* loci; and (ii) stable expression of the OsTir1 ligase into the *RCC1* locus. (C-E) Immunoblots showing degradation of LA/C (C), LB1 (D), and LB2 (E) compared to untagged DLD1 parental cell line after 16 hours of auxin treatment. (F) Quantification of nuclear envelope protein abundance by LC-MS/MS in nuclear extracts from mNG-AID-LA/C, mNG-AID-LB1, and mNG-AID-LB2 cells treated with auxin for 16 hours. (G) Number of genes significantly differentially expressed (*padj* < 0.05) at least 2-fold in mNG-AID-LA/C, mNG-AID-LB1, and mNG-AID-LB2 cells after 16 or 96 hours of auxin treatment compared to untreated mNG-AID-LA/C, mNG-AID-LB1, and mNG-AID-LB2 cells.

**Figure 2.**
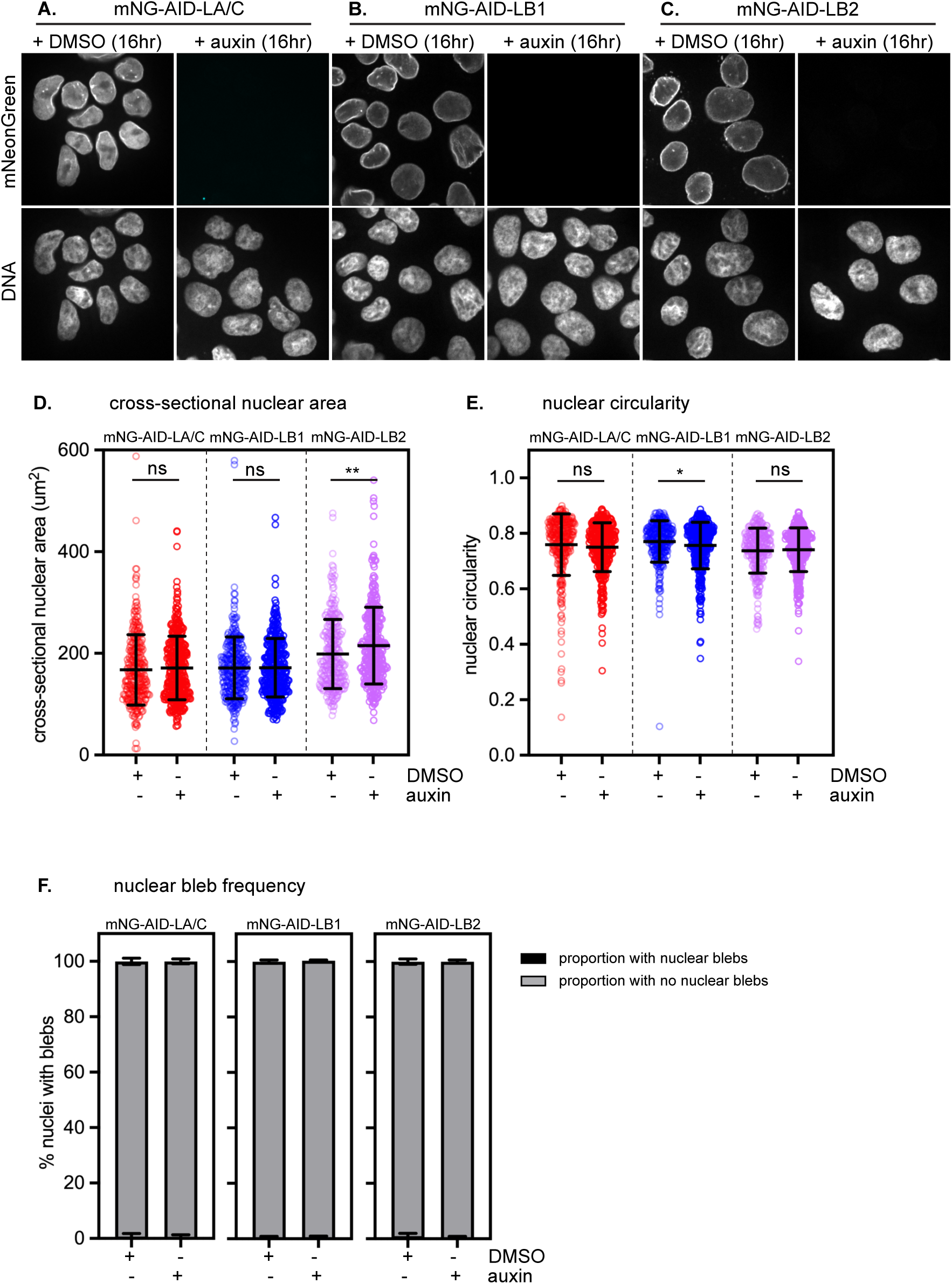
Acute laminate depletion does not drive changes in nuclear morphology. (A-C) Immunofluorescence of mNeonGreen after 16 hours of DMSO or auxin treatment in mNG-AID-LA/C (A), mNG-AID-LB1 (B), and mNG-AID-LB2 (C) cells. Scale bar, 10 μm. (D-E) Cross-sectional nuclear area (D) and nuclear circularity (E) after 16 hours of DMSO or auxin treatment in mNG-AID-LA/C, mNG-AID-LB1 and mNG-AID-LB2 cells. n > 275 cells analyzed per condition from 3 independent replicate experiments. ns indicates p > 0.05, ** indicates p < 0.01 and * indicates p < 0.05 by pairwise t-test. (F) Percentage of nuclei with blebs after 16 hours of DMSO or auxin treatment in mNG-AID-LA/C, mNG-AID-LB1, and mNG-AID-LB2 cells. n > 400 cells analyzed per condition from 3 independent replicate experiments.

With these degron cell lines in hand, we set out to evaluate the acute effects of lamin isoform depletion on nuclear organization and function in this cell type. DLD1 cells express high levels of the *LMNA* transcript and significantly lower levels of the B-type lamins (*LMNB1* and *LMNB2*) (Figure 1A). We first evaluated the acute consequences of depleting any lamin isoform on the nuclear enrichment of the other lamin isoforms or associated NE proteins. To accomplish this, we prepared lamina-enriched nuclear extracts after 12 hours of auxin treatment and quantified protein abundance by mass spectrometry (see Methods). Addition of auxin to mNG-AID-LA/C, mNG-AID-LB1 or mNG-AID-LB2 cells resulted in potent depletion of the targeted lamin isoform with no detectable changes in nuclear enrichment of the remaining isoforms (Figure 1F). Acute lamin isoform depletion had minimal effects on nuclear enrichment of lamina-associated proteins (for example, LBR, LUMA, LAP2α, or LAP2β; Figure 1F) or nuclear pore complex subunits (for example, Nup107, Nup160, and Nup205; Figure 1F). The Sun1 and Sun2 proteins appeared moderately sensitive to depletion of LB2 and LA/C, respectively (Figure 1F); this effect was reproducible for Sun1 but not Sun2 (Supplemental Figure 2A). Degradation of LA/C modestly but consistently decreased the nuclear enrichment of emerin (Figure 1F, Supplemental Figure 2A), consistent with emerin’s known reliance on LA/C for nuclear targeting (Sullivan *et al*., 1999). Altogether, these data indicate that acute lamin isoform depletion does not dramatically change the lamina-associated proteome, at least not within one cell cycle.

The nuclear lamina has been implicated in regulation of gene expression (Guelen *et al*., 2008; Kumaran and Spector, 2008). To query the immediate impact of lamin loss on gene expression, we performed RNAseq after acute depletion of LA/C, LB1, or LB2 for 16 or 96 hours (Fig 1G). To identify and control for the effects of auxin treatment on gene expression, we also performed RNAseq on DLD1 parental cells after treatment with DMSO vehicle or auxin for 16 or 96 hours. Any auxin-sensitive genes in lamin degron conditions were then excluded from further analysis (see Supplementary Table 2). We found that short (16 hour) depletion of each lamin isoform resulted in a relatively small number of differentially expressed genes (DEGs) (Figure 1G; Supplementary Table 3). After prolonged (96 hour) LA/C depletion, a larger number of DEGs accumulated (Figure 1G; Supplementary Table 3). In contrast, the number of DEGs after prolonged LB1 or LB2 depletion remained modest (Figure 1G; Supplementary Table 3). Overall, there was very little overlap in DEGs between lamin isoforms (Supplemental Figure 3B). These results indicate that dysregulation of gene expression is not a shared and immediate response to lamin depletion.

### Acute lamin depletion is not sufficient to drive changes in nuclear morphology

It has been previously reported that loss of any of the lamin isoforms can result in changes to nuclear morphology. For instance, KO or KD of LA/C decreases nuclear circularity (Sullivan *et al*., 1999; Broers *et al*., 2004; Lammerding *et al*., 2006; Robijns *et al*., 2016), while KO or KD of LB1 induces nuclear blebbing and/or micronucleus formation (Vergnes *et al*., 2004; Lammerding *et al*., 2006; Coffinier *et al*., 2011). We evaluated the acute effects of lamin depletion on nuclear morphology by quantifying nuclear cross-sectional area, nuclear circularity, and nuclear blebbing frequency in mNG-AID-LA/C, mNG-AID-LB1, and mNG-AID-LB2 cells after 16 hours of auxin treatment. Surprisingly, we generally failed to detect significant changes to nuclear shape or size in lamin-depleted nuclei (Figure 2A-E), except for a modest increase in nuclear cross-sectional area after LB2 depletion (Figure 2D) and a modest decrease in nuclear circularity after LB1 depletion (Figure 2E). Further, we failed to detect any increase to blebbing/micronucleation in lamin-depleted nuclei (Figure 2F). Altogether, these data indicate that acute lamin depletion is not sufficient to induce immediate nuclear morphology defects in this cell type.

### Lamin B1 depletion induces immediate lamin A/C meshwork disorganization

Each lamin isoform builds homotypic but overlapping polymeric meshworks at the nuclear periphery (Kittisopikul *et al*., 2019), and lamin filament assembly appears to be interdependent (Shimi *et al*., 2008; Guo *et al*., 2014). It has previously been reported that LB1 KD results in LA/C meshwork disorganization (Shimi *et al*., 2008). To determine whether LA/C meshwork disorganization is an immediate consequence of LB1 depletion, we treated mNG-AID-LB1 cells with auxin for 0, 2, 4, 8 and 16 hours and tracked LB1 abundance and LA/C organization by immunofluorescence microscopy. This analysis revealed that LA/C meshwork disorganization coincides with LB1 depletion, as each of these phenotypes become apparent within 4 hours of auxin treatment (Figure 3A-C). LB1 depletion is accompanied by opening of amorphous gaps in the LA/C meshwork, which are sometimes randomly distributed across the nuclear surface and are sometimes found at a nuclear pole (Figure 3A). LB1 depletion also induces similar disorganization of the INM protein lamin B receptor (LBR) (Supplemental Figure 4A-D), implying effects on both the lamina and other components of the nuclear periphery. Importantly, LB2 depletion had no effect on the LA/C meshwork or on LBR, indicating that these effects are specific to LB1 (Figure 3D-F, Supplemental Figure 4C-D) even though these homologous isoforms are expressed at comparable levels in DLD1 cells (Figure 1A).

**Figure 3.**
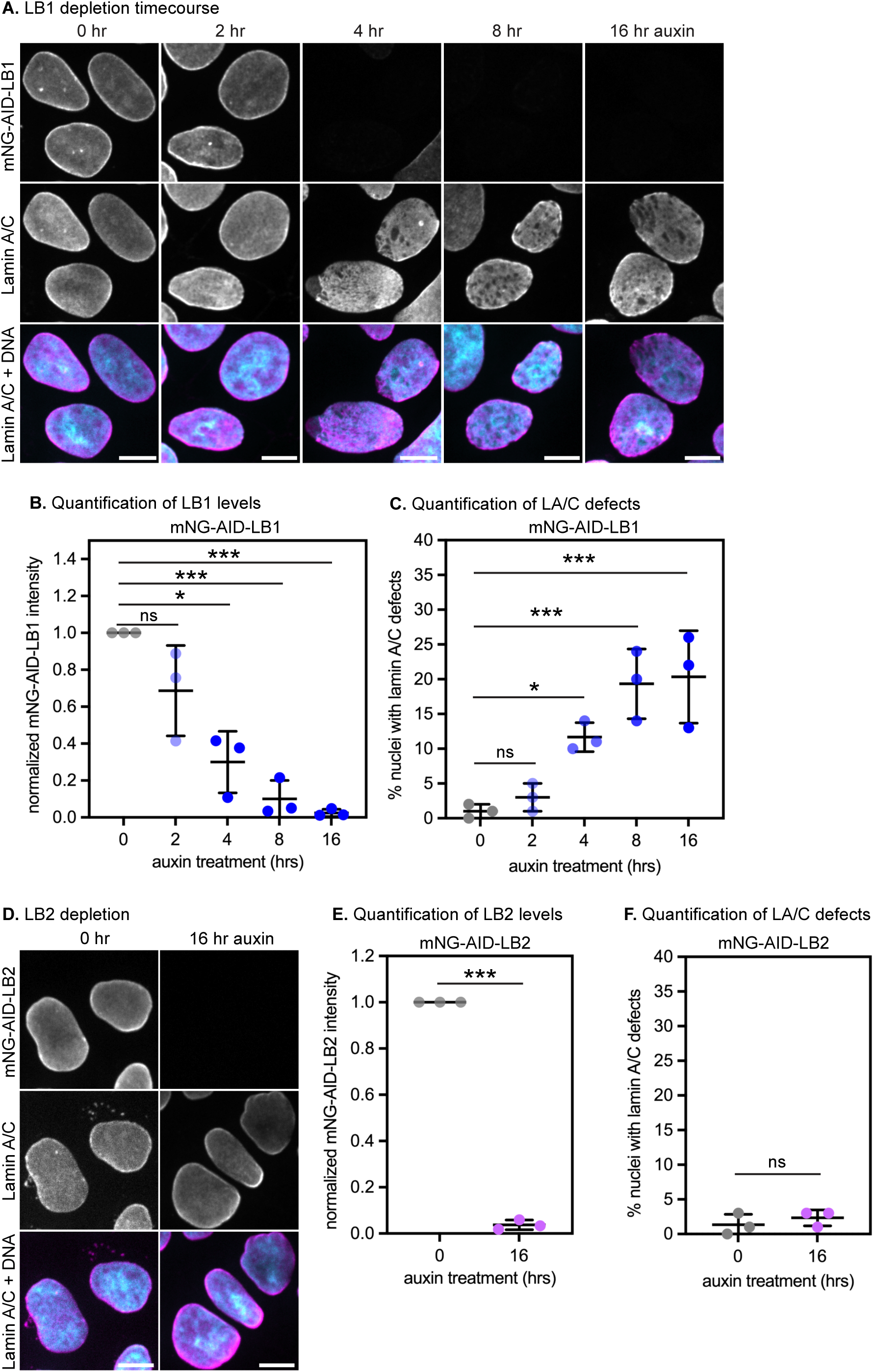
Lamin B1 depletion induces immediate lamin A/C meshwork disorganization (A) Immunofluorescence of mNeonGreen and lamin A/C after 0, 2, 4, 8, and 16 hours of auxin treatment in mNG-AID-LB1 cells. Scale bar, 10 μm (B) Normalized mNeonGreen intensity after 0, 2, 4, 8, and 16 hours of auxin treatment in mNG-AID-LB1 cells. (C) Percentage of nuclei with lamin A/C defects after 0, 2, 4, 8, and 16 hours of auxin treatment in mNG-AID-LB1 cells. In (B,C), ns indicates p > 0.05, * indicates p < 0.05, *** indicates p < 0.001 by one-way ANOVA followed by Dunnett’s multiple comparisons test. Average values of 3 independent replicate experiments shown; n > 350 cells analyzed per condition. (D) Immunofluorescence of mNeonGreen and lamin A/C after 16 hours of DMSO or auxin treatment in mNG-AID-LB2 cells. (E) Normalized mNeonGreen intensity after 16 hours of DMSO or auxin treatment in mNG-AID-LB2 cells. (F) Percentage of nuclei with lamin A/C defects after 16 hours of DMSO or auxin treatment in mNG-AID-LB2 cells. In (E,F), *** indicates p < 0.001 and ns indicates p > 0.05 by pairwise t-test. n > 300 cells analyzed per condition in 3 independent replicate experiments.

### Lamin A/C meshwork defects are reversed by actin depolymerization

Cytoskeletal forces are transmitted to the lamina via the <u>li</u>nker of <u>n</u>ucleoskeleton and <u>c</u>ytoskeleton (LINC) complex (King, 2023). Because LINC complex components interact with the A-type lamins (Crisp *et al*., 2005), we hypothesized that cytoskeletal forces on the LA/C meshwork induce defects in the absence of LB1. To test this hypothesis, we degraded LB1, then depolymerized the actin cytoskeleton by transient latrunculin A (Lat A) treatment (Figure 4A), which prevents actin polymerization by binding and sequestering monomeric actin (Coué *et al*., 1987) (Figure 4A). Lat A treatment increases nuclear height, confirming that the compressive effect of actin on the nucleus has been relieved (Figure 4B). Strikingly, 1 hour of actin depolymerization completely reverses the effects of LB1 depletion on LA/C meshwork organization (Figure 4C-G). This outcome is consistent with the model that LB1 promotes proper LA/C meshwork organization at least in part by resisting actin forces on the nucleus. The onset of LA/C meshwork defects upon LB1 loss and their resolution after actin depolymerization indicates that LA/C meshwork defects are rapid and reversible.

**Figure 4.**
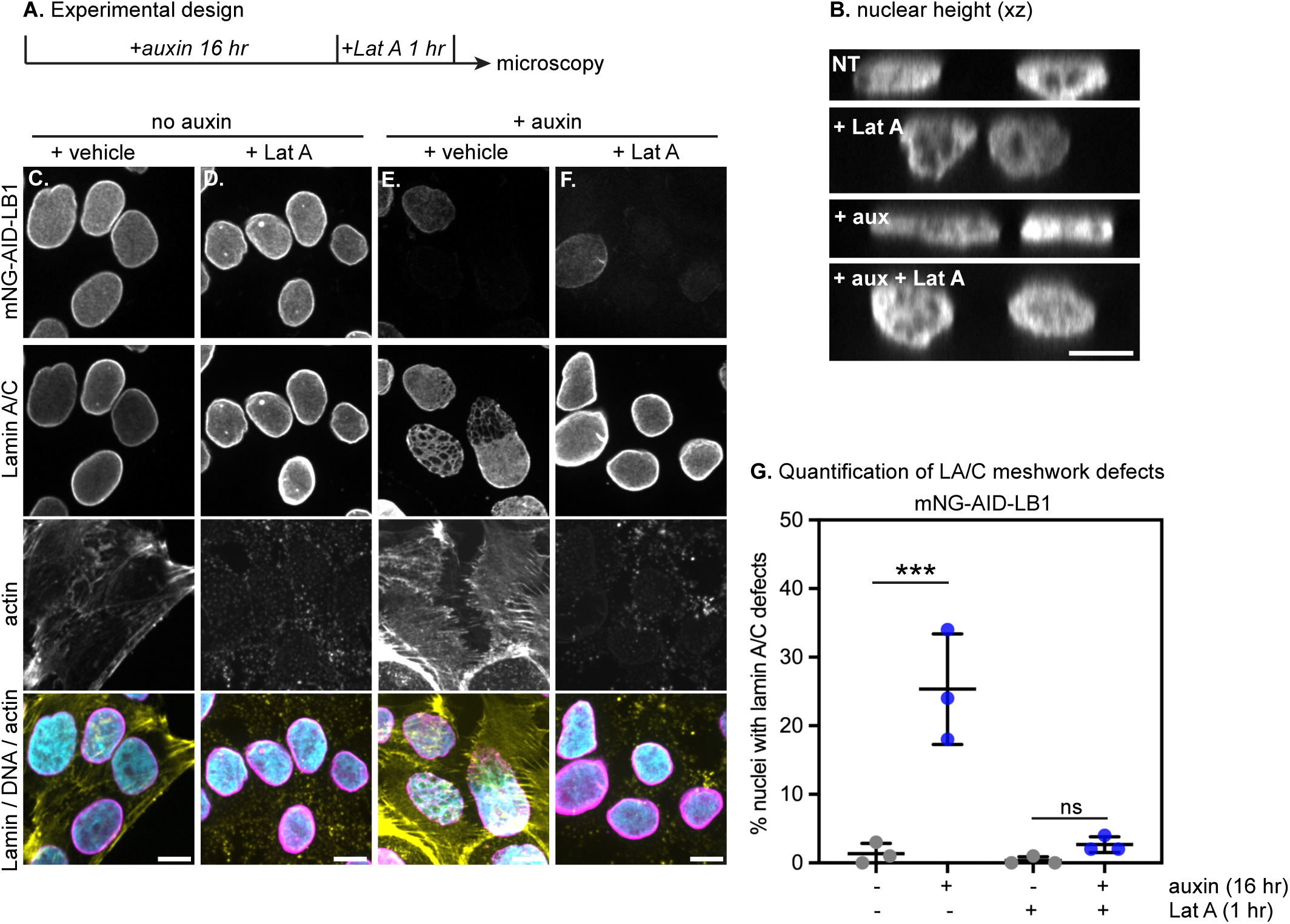
Lamin A/C meshwork defects are reversed by actin depolymerization (A) Overview of experimental strategy. mNG-AID-LB1 cells were treated with DMSO or auxin for 16 hours followed by an hour of DMSO or Latrunculin A treatment. mNG-AID-LB1 cells were then fixed and immunostained. (B) xz view to show nuclear height of mNG-AID-LB1 cells immunostained for lamin A/C after 16 hours of DMSO or auxin treatment followed by 1 hour of additional DMSO or Latrunculin A. (C-F) Immunofluorescence of mNeonGreen, lamin A/C and actin after mNG-AID-LB1 cells were treated with (C) DMSO or (E) auxin for 16 hours. mNG-AID-LB1 cells were then treated with (D) Latrunculin A for an hour or (F) 16 hours of auxin followed by 1 hour of Latrunculin A treatment. Scale bar, 10 μm (G) Quantification of the percentage of nuclei with lamin A/C defects after mNG-AID-LB1 cells were treated with DMSO or auxin for 16 hours followed by 1 hour of additional DMSO or Latrunculin A treatment. n > 400 cells analyzed per condition in 3 independent replicate experiments.

### Neither Lamin A/C nor Lamin B1 depletion are sufficient to cause nuclear rupture

Our data indicate that LB1 depletion rapidly induces defects in the LA/C meshwork (Figure 3A, C). Gaps in the lamin meshwork are thought to weaken the nucleus and predispose nuclei to rupture (Maciejowski and Hatch, 2020). To determine whether acute LB1 and/or LA/C depletion induce nuclear rupture, we generated dual color nuclear rupture reporter cell lines stably expressing the tdTomato fluorescent protein fused to 3 tandem nuclear localization signals (3xNLS-tdTomato) and histone H2B fused to a monomeric near-infrared fluorescent protein (H2B-mIRFP). In this system, the presence of cytosolic 3xNLS-tdTomato indicates nuclear rupture while H2B-mIRFP marks nuclei. We imaged mNG-AID-LA/C and mNG-AID-LB1 nuclear rupture reporter cells in the absence or presence of auxin at 3-minute intervals for 4 hours (Figure 5; Supplementary Movie 1,2). Surprisingly, we observed no significant increase in nuclear rupture in the absence of either LA/C (Figure 5A-C) or LB1 (Figure 5D-F). These data indicate that LB1 depletion and subsequent LA/C meshwork disorganization are not sufficient to cause nuclear rupture in this cell type.

**Figure 5.**
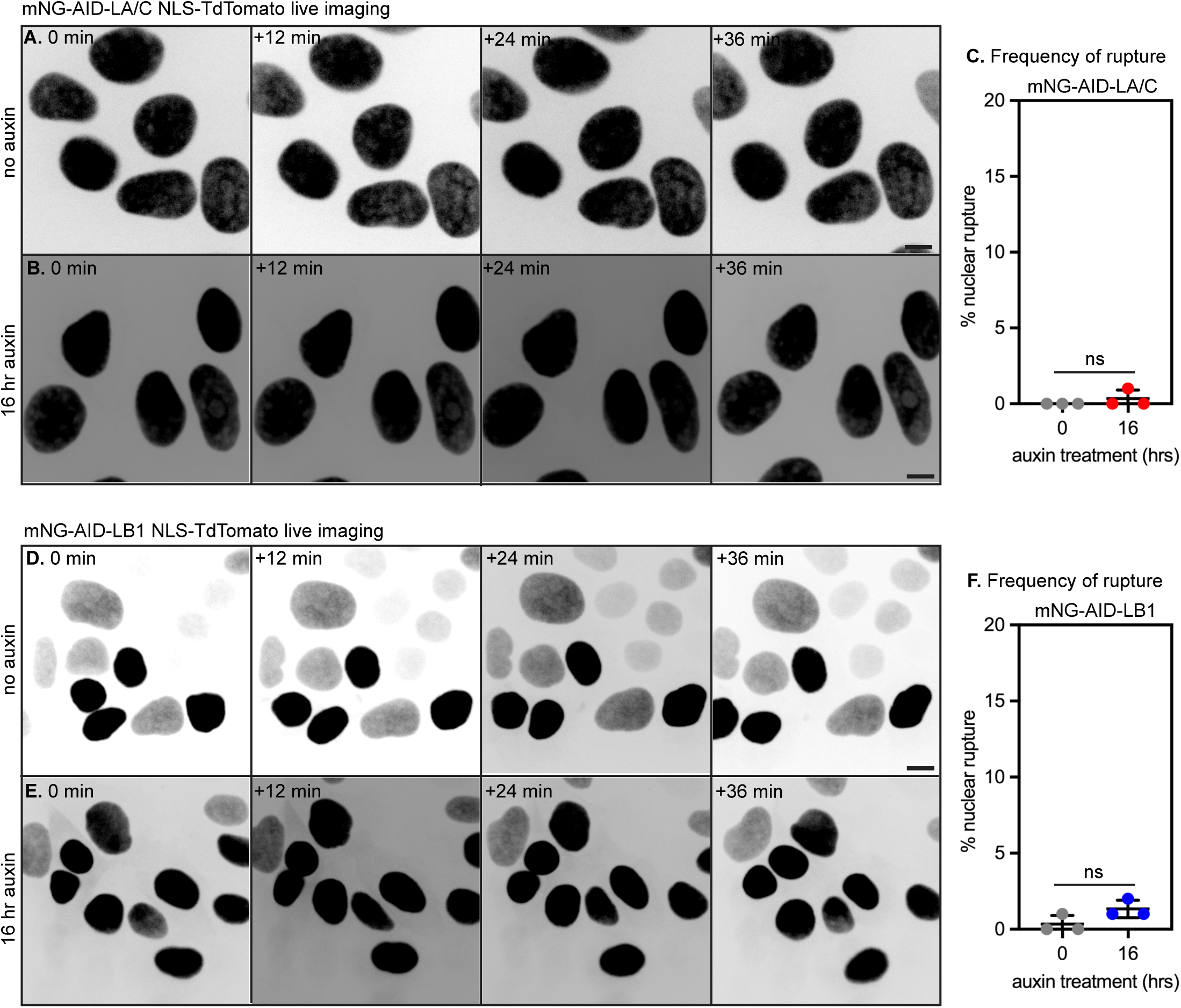
Neither Lamin A/C nor Lamin B1 depletion are sufficient to cause nuclear rupture (A-B) Still images from time-lapse imaging of mNG-AID-LA/C cells stably expressing 3xNLS-tdTomato and treated with DMSO vehicle control (A) or with auxin (B) for 16 hours. (C) Quantification of NE rupture in mNG-AID-LA/C cells after 16 hours of DMSO or auxin treatment. Average values for replicate experiments shown; n > 400 cells analyzed across 3 replicate experiments. (D-E) Still images from time-lapse imaging of mNG-AID-LB1 cells expressing 3xNLS-tdTomato and treated with DMSO vehicle control (D) or with auxin (E) for 16 hours. Scale bar, 10 μm. (F) Quantification of NE rupture in mNG-AID-LB1 cells after 16 hours of DMSO or auxin treatment. Average values for replicate experiments shown; n > 600 cells analyzed across 3 replicate experiments.

### LAP2β and Lamin B1 co-depletion increases lamin A/C meshwork defects

The lamins are thought to play a major role in maintaining nuclear integrity, and nuclei harboring lamin mutations or lacking lamin isoforms have been shown to be more prone to rupture (Vos *et al*., 2011; Vargas *et al*., 2012; Tamiello *et al*., 2013; Comaills *et al*., 2016; Hatch and Hetzer, 2016; Chen *et al*., 2018; Xia *et al*., 2018; Chen *et al*., 2019). Considering this precedent, we were surprised that we did not detect nuclear rupture when either LA/C or LB1 were acutely depleted (Figure 5). We hypothesized that lamin-interacting INM proteins might maintain nuclear integrity in the absence of LA/C or LB1. We evaluated the expression levels of INM proteins in DLD1 cells by RNAseq, which revealed that the LEM family protein LAP2β (Barton *et al*., 2015), encoded by the *TMPO* gene, was very highly expressed - more highly even than the *LMNB1* and *LMNB2* transcripts (Fig 6A). This observation aligns with a previous report that LAP2β is highly expressed in gastrointestinal cancers (Kim *et al*., 2012), as DLD1 cells are of colon cancer origin. LAP2β has been recently shown to counteract progerin-induced lamin meshwork disorganization and to protect LB1-null neurons from nuclear rupture (Chen *et al*., 2021; Kim *et al*., 2024). These findings led us to hypothesize that high levels of LAP2β protect lamin-depleted DLD1 cells from nuclear rupture.

**Figure 6.**
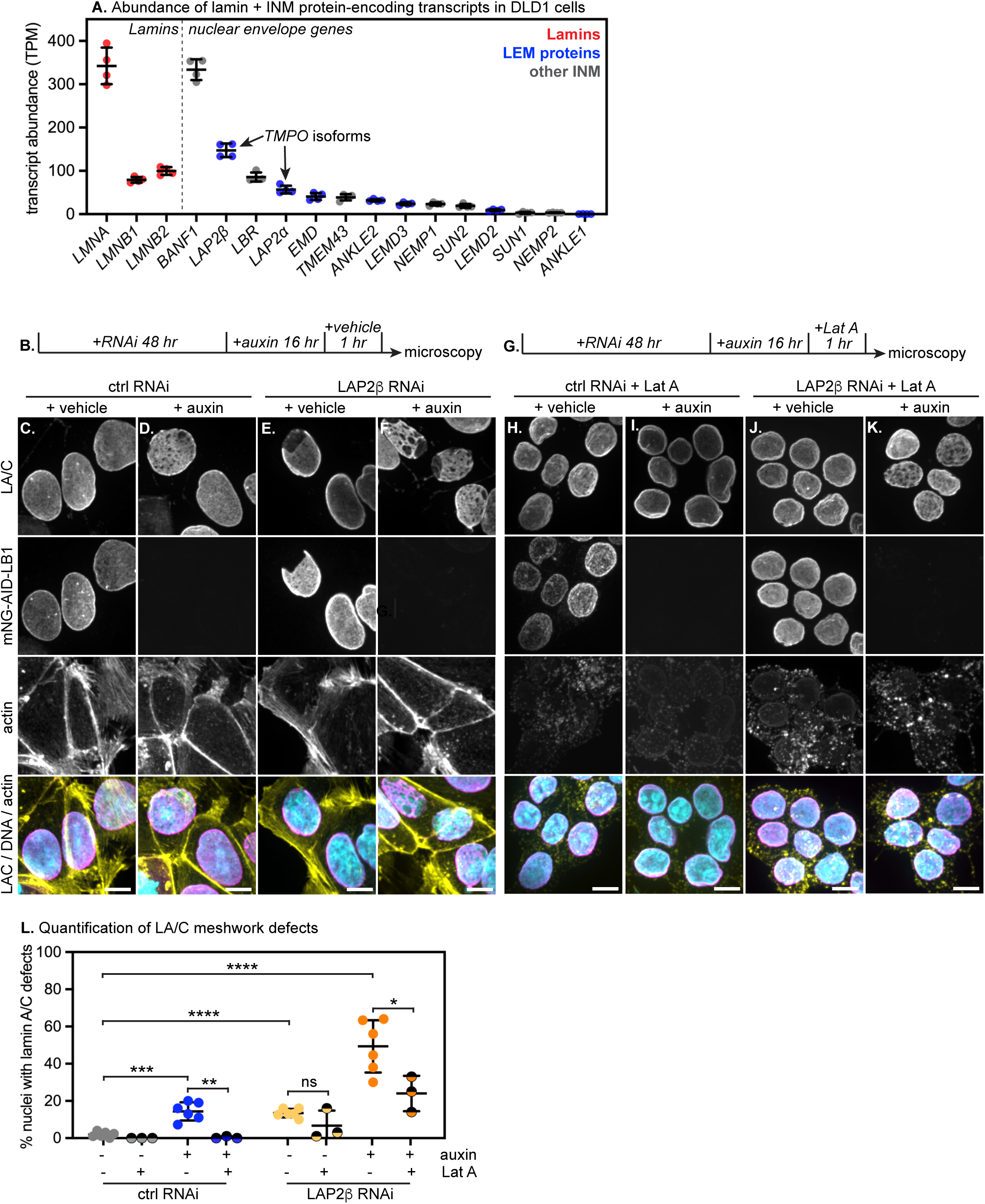
LAP2β and Lamin B1 co-depletion increases lamin A/C meshwork defects in an actin-dependent manner (A) Relative transcript abundance of lamins and various INM proteins in DLD1 cells; Nucleoplasmic *TMPO* splice isoform LAP2α distinguished from INM-localized splice isoform LAP2β by isoform-resolved analysis with Salmon (see Methods). (B) Overview of experimental strategy. mNG-AID-LB1 cells were subject to RNAi for 48 hours followed by 16 hours of auxin treatment. (C-F) Immunofluorescence of lamin A/C, mNG-AID-LB1, and actin (phalloidin) in mNG-AID-LB1 cells after control RNAi (C-D) or LAP2β RNAi (E-F) along with vehicle (C,E) or auxin (D,F). All conditions received 1 hour vehicle treatment as control for G-K. (G) Overview of experimental strategy. mNG-AID-LB1 cells were subject to RNAi for 48 hours followed by 16 hours of auxin treatment, then 1 hour of Lat A treatment. (H-K) Immunofluorescence of lamin A/C, mNG-AID-LB1, and actin (phalloidin) in mNG-AID-LB1 cells after control RNAi (H-I) or LAP2β RNAi (J-K) along with vehicle (H,J) or auxin (I,K). All conditions received 1 hour Lat A treatment. Scale bar, 10 μm. (L) Quantification of the percentage of nuclei with lamin A/C defects. Each dot on graph represents one experimental replicate; n > 400 cells analyzed per condition across at least 3 replicate experiments. ns indicates p > 0.05, * indicates p < 0.05, ** indicates p < 0.01, *** indicates p < 0.001, and **** indicates p < 0.0001 by unpaired t-test.

To test this hypothesis, mNG-AID-LB1 cells were subjected to small interfering RNA (siRNA)-mediated knockdown of LAP2β using validated isoform-specific siRNAs (Mirza *et al*., 2019) (Figure 6B; Supplemental Figure 5A-C). We found that LAP2β knockdown caused defects in both LA/C and LB1 meshwork organization (Figure 6E,L, Supplemental Figure 5C). We could visualize the dynamics of LB1 meshwork defects over time in living LAP2β-depleted mNG-AID-LB1 cells by imaging the mNG tag (Supplementary Movie 3). LB1 depletion along with LAP2β knockdown further disorganized the LA/C meshwork, inducing both larger and more frequent LA/C meshwork defects (Figure 6F,L). LA/C meshwork defects caused by LAP2β depletion were improved by Lat A-mediated actin depolymerization (Figure 6G-L). To confirm these results, we targeted LAP2β with RNAi in DLD1 cells (Supplemental Figure 6A-B). In contrast to our findings in the mNG-AID-LB1 background, knockdown of LAP2β alone in DLD1 cells did not induce LA/C defects. However, knockdown of both LB1 and LAP2β together did (Supplemental Figure 6A-B). This outcome indicates that mNG-AID-LB1 cells are more sensitive to LAP2β depletion than normal DLD1 cells. We infer that the expression and/or function of LB1 is weakened by the introduction of the mNG-AID tag in the mNG-AID-LB1 cell line. We noted that tagging moderately reduces LB1 transcript and protein expression (Supplemental Figure 3C; Figure 1D). As endogenous N-terminal tagging interferes with the function of LA/C (Odell and Lammerding, 2024), it is also possible that the mNG-AID tag impairs LB1 function. We therefore conclude that disruption of LB1 and LAP2β together causes LA/C meshwork defects and infer that these proteins work synergistically to promote the normal organization of the NL.

### LAP2β and Lamin B1 co-depletion induces nuclear rupture

Co-depletion of LAP2β and LB1 induced both larger and more frequent LA/C defects than either perturbation alone (Figure 6F,L), leading us to ask whether these perturbations would together cause nuclear rupture. To evaluate the effect of LAP2β loss on nuclear integrity, we knocked down LAP2β and performed live imaging in mNG-AID-LB1 cells stably expressing tdTomato-3xNLS and H2B-mIRFP. LAP2β knockdown caused a low but significant incidence of nuclear rupture in mNG-AID-LB1 cells (Figure 7D,I; Supplementary Movie 4), while depletion of LB1 alone did not induce significant levels of nuclear rupture (Figure 5D-F; Figure 7C,I). The frequency of rupture was significantly increased by the co-depletion of both LAP2β and LB1 (Figure 7E,I; Supplementary Movie 5). This result indicates that LAP2β expression can protect nuclear integrity in the absence of LB1, consistent with recent studies in LB1 KO cells (Chen *et al*., 2021).

**Figure 7.**
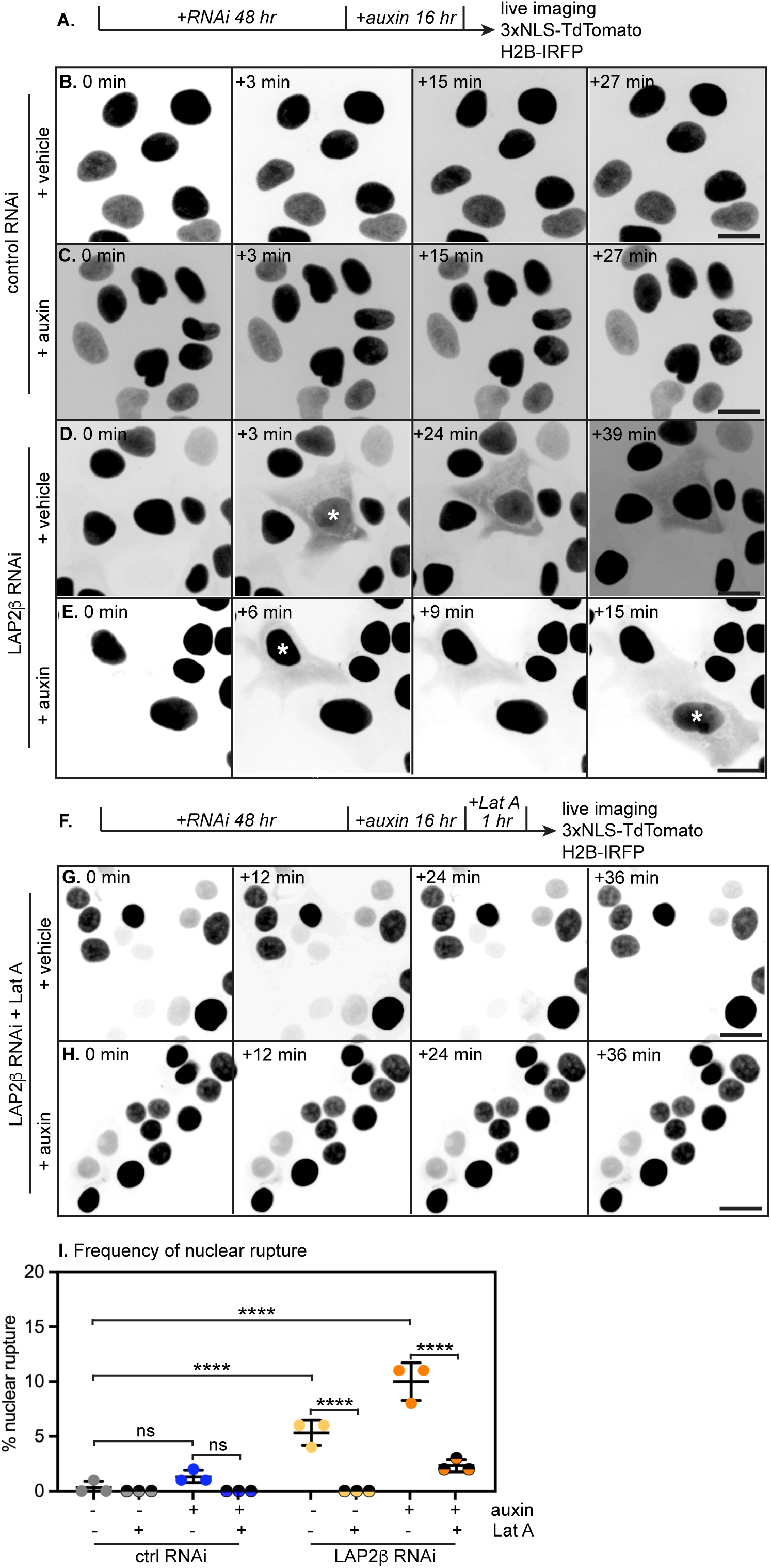
LAP2β and Lamin B1 co-depletion induces nuclear rupture in an actin-dependent manner (A) Overview of experimental strategy. mNG-AID-LB1 cells were subject to RNAi for 48 hours followed by 16 hours of DMSO or auxin treatment and live imaging at 40x magnification with images of 3xNLS-TdTomato and H2B-IRFP acquired every 3 minutes for 4 hours. (B-E) Still images from time-lapse imaging of mNG-AID-LB1 cells expressing 3xNLS-tdTomato and treated with control RNAi (B), control RNAi + auxin(C), LAP2β RNAi (D), or LAP2β RNAi + auxin (E). See also Supplementary Movies 4 and 5. (F) Overview of experimental strategy. mNG-AID-LB1 cells were subject to RNAi for 48 hours followed by 16 hours of DMSO or auxin treatment and 1 hour of Lat A treatment, followed by live imaging at 40x magnification with images of 3xNLS-TdTomato and H2B-mIRFP acquired every 3 minutes for 4 hours. (G-H) Still images from time-lapse imaging of mNG-AID-LB1 cells expressing 3xNLS-tdTomato and treated with LAP2β RNAi and Lat A (G) or LAP2β RNAi + auxin + Lat A (H). See also Supplementary Movies 7 and 8. (I) Quantification of NE rupture in AID-Lamin B1 cells after indicated RNAi, 16 hours of DMSO or auxin followed by additional DMSO or Lat A. Each dot on graph represents one replicate; n > 200 cells analyzed per condition. Scale bar, 20 μm. ns indicates p > 0.05, **** indicates *padj* < 0.0001 by unpaired t-test.

The compression of the nucleus by the actin cytoskeleton has previously been shown to drive nuclear rupture (Hatch and Hetzer, 2016). We wanted to determine whether actin forces contribute to nuclear rupture in DLD1 cells depleted of LAP2β and LB1. To test this possibility, we depleted LAP2β and/or LB1, then treated cells with Lat A for 1 hour before live imaging. Actin depolymerization strongly reduced nuclear rupture frequency when LAP2β or both LAP2β and LB1 were depleted (Figure 7F-I; Supplementary Movie 7,8). We conclude that nuclear rupture induced by LAP2β knockdown and lamin B1 is caused by the compressive force of actin.

We could compare lamina defects to sites of nuclear ruptures in LAP2β-depleted cells by imaging mNG-AID-LB1 and the 3xNLS-TdTomato reporter, respectively (Supplementary Movie 6). These analyses suggest that nuclear ruptures occur at membrane sites that overlay large lamina defects (Figure 8, Supplementary Movie 6). We also found examples of ruptures that were preceded by localized nuclear blebbing within large lamina defects (Figure 8, yellow arrowheads; Supplementary Movie 6). Interestingly, these observations indicate that lamina defects do not induce the entire overlying membrane area to bleb, but rather that blebbing initiates at a localized membrane site.

**Figure 8.**
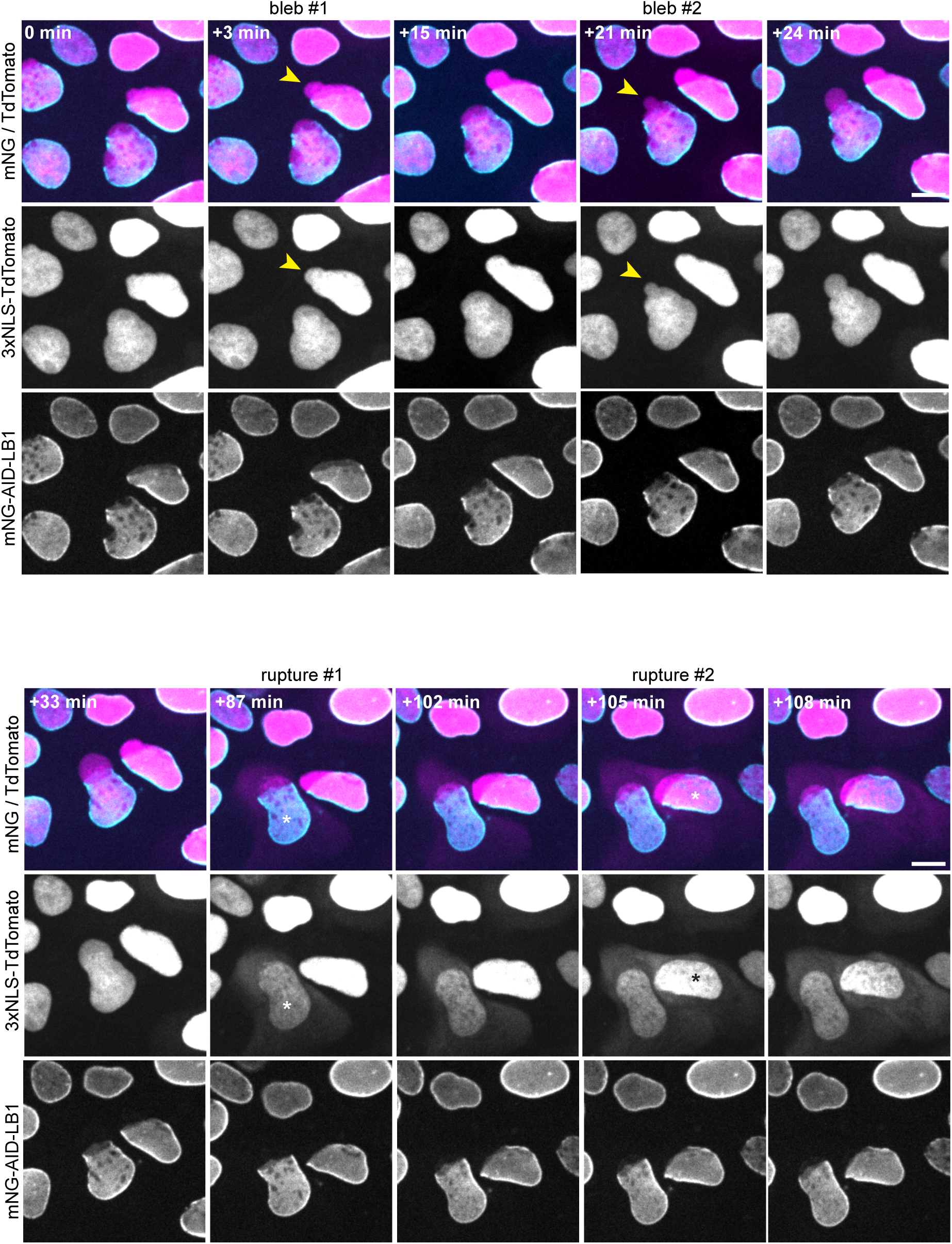
Nuclear ruptures occur within lamina gaps in LAP2β-depleted mNG-AID-LB1 cells. Still images from time-lapse imaging of mNG-AID-LB1 cells treated with LAP2β RNAi and expressing 3xNLS-TdTomato and H2B-IRFP (channel not shown). Yellow arrowheads indicate nuclear blebs and * indicates nuclear ruptures. Scale bar, 10 μm. See also Supplementary Movie 6.

## Discussion

In this study, we took advantage of the auxin-inducible degron system to dissect the immediate impact of lamin loss on nuclear function. We explored the influence of the lamins on the composition of the NE proteome (Figure 1F), gene expression (Figure 1G), nuclear morphology (Figure 2), and nuclear integrity (Figure 5). Altogether, these studies indicated significant latency and/or pleiotropy in several phenotypes that have previously been associated with loss of lamin function.

The NL has been linked to regulation of gene expression (Briand and Collas, 2020). However, we found that acute depletion of each lamin isoform (16 hours) had little effect on gene expression (Figure 1G), indicating that gene expression changes are not an immediate consequence of lamin depletion. Prolonged depletion (96 hours) of LA/C did induce more widespread dysregulation of gene expression (Figure 1G). When incorporated into the NL, LA/C has been implicated in repression (Solovei *et al*., 2013), while nucleoplasmic LA/C, in contrast, has been associated with active transcription (Gesson *et al*., 2016; Ikegami *et al*., 2020). Interestingly, a nucleoplasmic pool of mNG-AID-LA/C is readily detectable in DLD1 cells (Figure 2A), and we observed many downregulated genes after prolonged LA/C depletion (Figure 1F), implying that LA/C may contribute to active transcription in DLD1 cells.

Many prior studies have linked lamin perturbations to nuclear deformations in primary, immortalized, and transformed cell lines (Vergnes *et al*., 2004; Lammerding *et al*., 2006; Coffinier *et al*., 2011; Schibler *et al*., 2023). We did not detect such changes in nuclear shape after acute depletion of any lamin isoform in DLD1 cells (Figure 2). This outcome indicates that acute lamin isoform depletion is not sufficient to induce either nuclear shape changes or nuclear blebbing. There are several potential explanations for the inconsistency between our findings and previous reports. It is possible that nuclear shape changes are a secondary consequence of lamin isoform loss that become apparent only after multiple cell cycles; however, we did not observe any increase in nuclear deformations or blebbing after prolonged (96 hours) auxin treatment (not shown). Alternatively, it is possible that the influence of orthogonal regulators of nuclear morphology varies across cell types. For instance, heterochromatin compaction and actin contractility each influence nuclear blebbing propensity (Stephens *et al*., 2018; Pho *et al*., 2024) and are variable across cell types. Notably, a recent RNAi screen for regulators of nuclear shape found extensive variation between two cell lines (Schibler *et al*., 2023). Depletion of various chromatin regulators influenced nuclear shape, but one of the few consistent hits in both cell types was LA/C (Schibler *et al*., 2023). It has been established that the relative stoichiometry of A-type to B-type lamins influences NL properties (Swift *et al*., 2013). Given the high expression of A-type lamins in DLD1 cells (Figure 1A), it is surprising that acute LA/C depletion had no detectable effect on nuclear morphology (Figure 2) or on nuclear integrity (Figure 5A-C).

We determined that LA/C meshwork defects are an immediate consequence of LB1 but not LB2 depletion (Figure 3), indicating that this previously described phenomenon (Shimi *et al*., 2008; Hatch and Hetzer, 2016) is a rapid, specific, and cell type-invariant consequence of LB1 loss. LA/C defects are rapidly and potently corrected by Lat A-mediated actin depolymerization (Figure 4). This outcome implies that gaps in the LA/C meshwork are a result of increased vulnerability to actin forces in the absence of LB1. On average, approximately 20% to 30% of nuclei experience LA/C meshwork disorganization because of LB1 depletion. One would expect that prolonged periods of LB1 depletion would increase the number of nuclei with LA/C defects, however we did not see this even after prolonged (96 hours) auxin treatment (not shown). We speculate that there are other factors (such as LAP2β) involved in LA/C meshwork organization that keep the lamina intact when LB1 is depleted. Additionally, it could be that the cells that do show LA/C disorganization when LB1 is depleted experience greater effects of actin.

Nuclear rupture is thought to occur at NL gaps (Maciejowski and Hatch, 2020). However, we found that LB1 depletion caused LA/C meshwork defects without inducing nuclear rupture in DLD1 cells (Figure 5D-F). This observation indicates that nuclear integrity can be maintained when gaps in the NL are present, and hints that additional factors influence nuclear integrity. We identified the highly expressed INM protein LAP2β as one such factor (Figure 6-7). LAP2β and LB1 co-depletion increased the size and frequency of NL gaps (Figure 6B-F) and induced nuclear ruptures (Figure 7D-E).

Several lines of evidence suggest that LAP2β has a major influence on NL function. LAP2β overexpression inhibits nuclear rupture in LB1-depleted cells (Maciejowski *et al*., 2015; Chen *et al*., 2021) and can even mitigate the dominant, toxic effects of the mutant LA/C isoform progerin on NL organization and nuclear integrity (Kim *et al*., 2024). Our findings add to this body of evidence by demonstrating that LB1 and LAP2β co-depletion induces major NL defects. LAP2β binds LB1 but not LA/C (Foisner and Gerace, 1993). In addition to binding LB1, LAP2β interacts with the DNA-crosslinking protein barrier to autointegration factor 1 (BANF1/BAF) via a conserved LEM domain and binds directly to DNA via a divergent LEM-like domain (Cai *et al*., 2001). While BAF and other LEM proteins participate in the repair of nuclear ruptures, LAP2β has not been implicated in this process; it is not recruited to sites of rupture (Halfmann *et al*., 2019) and its depletion does not prolong nuclear rupture(Chen *et al*., 2021). Altogether, these pieces of evidence suggest that LAP2β could enhance nuclear integrity by tethering the INM to both LB1 and chromatin. The roles of LAP2β’s protein:protein and protein:DNA interactions in maintaining nuclear integrity will require future investigation.

## Materials and Methods

### Degron cell line generation

CRISPR/Cas9-mediated homologous recombination was used to introduce tags into the 5’ end of each lamin isoform locus (*LMNA, LMNB1,* and *LMNB2).* Cas9 was targeted to these loci by gRNAs cloned into the pX330 vector backbone (Addgene #42230); gRNA sequences are listed in Supplementary Table 3. For each target locus, an HR template was cloned including 5’ and 3’ homology arms amplified from genomic DNA extracted from DLD1 cells, except 3’ homology arm for *LMNB2*, which was synthesized by Synbio Technologies. These homology arms flank a hygromycin resistance gene, P2A sequence, mNeonGreen tag (Aksenova *et al*., 2020), and micro-AID tag (Morawska and Ulrich, 2013b).

After AID tag insertion, cells were subjected to a second round of CRISPR/Cas9 editing to introduce P2A-9xMyc-Tir1 into the 3’end of the *RCC1* locus. Lamin locus edits were verified by genotyping PCR for homozygous insertion of tags with primers listed in Supplementary Table 4.

### Cell culture

DLD1s were cultured in Dulbecco’s modified Eagle’s medium (DMEM; GenClone 25-501) supplemented with 10% fetal bovine serum. Cells were incubated at 5% CO2 and at 37 °C. Auxin (Sigma Aldrich I5148) was dissolved in DMSO (where indicated) or water and used at a concentration of 1mM. Latrunculin A (Thermo Scientific L12370) was dissolved in DMSO and used at a concentration of 1 µm.

### Analysis of nuclear extracts by mass spectrometry

A lamina / NPC-enriched fraction was isolated according to Cronshaw (Cronshaw *et al*., 2002), with modifications as recently described (Aksenova *et al*., 2020). Cells were incubated for 2 x 5 minutes in buffer A (20 mM HEPES, pH 7.8; 2 mM DTT, 10% sucrose, 5 mM MgCl2, 5 mM EGTA, 1% Tx100, 0.075% SDS) at room temperature, followed by incubation in buffer B (20 mM HEPES, pH 7.8; 2 mM DTT, 10% sucrose, 0,1 mM MgCl2), containing 4 ug/ml RNAse A (Promega) for 10 min at 37°C. Cells were then washed in buffer A. Cells were then incubated in buffer C (20 mM HEPES, pH 7.8; 150 mM NaCl, 2 mM DTT, 10% sucrose, 0.3% Empigen BB) for 10 min at 37 °C. The supernatant was spun for 3 min at 28,000 x g at 4 °C, then transferred to fresh tubes. Saturated TCA was added to a final concentration of 8%, followed by centrifugation and washing the pellet in ethanol. The pellet was finally solubilized in buffer D (8 M urea, 5 mM DTT) by pipetting and vortexing, then centrifuged for 1 min at 11,000 x g. The supernatant was collected for protein concentration and proteomic analysis.

Samples were analyzed by liquid chromatography mass spectrometry (LC-MS) in label-free quantitation mode (LFQ, Figure 1F) or as multiplexed tandem mass tagged (TMT) samples (Supplemental Figure 2A). In the latter case, samples were labeled with TMT reagents (TMT10plex label reagent set, Thermo Scientific) according to the manufacturer’s instructions. Samples were run on an LTQ Orbitrap Lumos nanoLCMS system (Thermo Fisher Scientific). Protein identification and quantification was performed in Proteome Discoverer (Thermo Fisher Scientific). Peptide IDs were assigned by searching the raw data agains the UniProt human database using the Mascot algorithm (Matrix Science Inc); peptide-spectrum matches were further validated with the Percolate algorithm. The median values of peptide-level fold changes were calculated to determine protein-level fold changes.

### siRNAs

siRNAs specifically against LAP2β (Mirza *et al*., 2019) were ordered from Sigma-Aldrich with the following sequences: LAP2β.1, 5′-CAGAGGAGCGAAGAGUAGA[dT][dT]-3′ LAP2β.2, 5′-GCAUGCAUCUUCUAUUCUG[dT][dT]-3′ and LAP2β.3, 5′-UGGACGAGUGGAUCUUCAA[dT][dT]-3′. MISSION® esiRNA against LMNB1 were ordered from Sigma-Aldrich (EHU057911). A non-targeting control siRNA was ordered from Dharmcon (D-001810-01-20). Cells were transfected with 125 nM RNAi using Lipofectamine 2000 (Life Technologies 11668019) transfection reagent according to the manufacturer’s instructions. Cells were incubated for at least 48 hours after RNAi transfection.

### Immunofluorescence

Cells in Ibidi chambers (Ibidi USA 80826) were fixed in 4% PFA for 5 min, and permeabilized and blocked in PBS, 0.1% Tx100, 0.02% SDS, 10 mg/ml BSA (IF buffer) before staining with primary and secondary antibodies diluted in IF buffer. Primary antibody dilutions were as follows: Lamin A/C (1:1000, Santa Cruz Biotechnology sc-376248), Lamin B1 (1:500, Santa Cruz Biotechnology sc-365214), Lamin B2 (1:500, Thermo Scientific 33-2100), LAP2β (1:400, Thermo Scientific PA5-52519), LBR (1:1000, abcam ab232731), and NPCs (1:1000, abcam ab24609). For phalloidin staining, cells in Ibidi chambers were fixed in 3.7% PFA for 10 min, permeabilized in 0.5% Triton X-100 in PBS for 5 min and blocked in 1mg/ml BSA for an hour. Cells were incubated with conjugated 568-Phalloidin (1:400, Thermo Scientific A12380) in 1mg/ml BSA in PBS.

### Nuclear rupture plasmid and cell line generation

The nuclear rupture reporter plasmid was generated by cloning H2B-mIRFP (Piatkevich *et al*., 2018) and 3(SV40)NLS-tdtomato (Adam *et al*., 1989; Shaner *et al*., 2004) separated by a P2A (Kim *et al*., 2011) sequence into a PiggyBac plasmid via Gibson assembly. To generate stable cell lines, mNG-AID-LA/C, mNG-AID-LB1, and mNG-AID-LB2 cells were transfected with the nuclear rupture reporter plasmid and PiggyBac transposase plasmid using jetprime transfection reagent (Polyplus 101000027). AID cells were selected in blasticidin (Thermo Fisher Scientific A1113902) for 7 days.

### Image Acquisition and analysis

Images were taken using an inverted Nikon Ti microscope equipped with a CSU-X1 spinning disk confocal with a 60 x 1.4 NA oil objective. Images are shown as maximum intensity projections of z stacks. Nuclear area was quantified by measuring the cross-sectional area occupied by a mask corresponding to a DNA stain. Lamin B1 signal intensity was quantified by measuring the maximum intensity projections of the Lamin B1 channel by a mask corresponding to a DNA stain. Nuclear shape was determined based off a mask of the DNA stain. All image quantification was performed with ImageJ. LA/C defects and nuclear blebbing defects were quantified by blind scoring.

### Time lapse imaging and nuclear rupture analysis

Time lapse images were taken using an inverted Nikon Ti microscope equipped with a CSU-X1 spinning disk confocal and an OkoLab cage incubator with CO2 and humidity control with a 40 x 0.95 NA objective. Cells in ibidi chambers were kept at 5% CO2 and 37 °C. Images were acquired every 3 min for 4 hours. For actin depolymerization experiments, Latrunculin A was added 1 hour before the start of image acquisition. To quantify nuclear rupture, all nuclei in the field were counted then the presence of H2B-mIRFP marked nuclei and cytoplasmic tdTomato were manually counted and considered a rupture event.

### Western blot assay

Cell pellets were lysed with a buffer containing 8M Urea, 75mM NaCl, 50mM Tris pH 8.0, and one tablet of cOmplete, Mini, EDTA-free Protease Inhibitor Cocktail (Roche 11836170001). Protein lysates were quantified using Pierce™ BCA Protein Assay Kits (Thermo Scientific 23225). 20 µg of protein was loaded per well. Protein lysates were run on mini-PROTEAN precast 4-15% gels (BioRad 4561094) followed by wet transfer onto 0.45 µm nitrocellulose membranes (GVS 1212590). Blots were probed with Lamin A/C (1:1000, Active Motif 39288), Lamin B1 (1:500, Santa Cruz Biotechnology sc-365214), Lamin B2 (1:500, Thermo Scientific 33-2100), and GAPDH (1:1000, GeneTex GTX637966) primary antibodies in 5% milk TBST. Anti-mouse HRP-conjugated (1:5000, Rockland 610-1302) and anti-rabbit HRP-conjugated (1:5000, Rockland 611-130) secondary antibodies were used. Blots were visualized using Pierce ECL Western Blotting Substrate (Thermo Scientific 32209) and Radiance Plus – Chemiluminescent Substrate (Azure Biosystems AC2103).

### Statistical analysis

GraphPad Prism9 was used to generate all graphs and perform statistical analyses.

### RNAseq library preparation

DLD1 parental, mNG-AID-LA/C, mNG-AID-LB1, and mNG-AID-LB2 cells were treated with auxin for 0, 16 or 96 hours. 3-4 replicates per condition were used. Each replicate was one well of a 6-well plate (Corning 3516). RNA was extracted using RNeasy Plus Mini Kit (QIAGEN 74136). Before prepping the libraries, the RNA was DNAse treated using the TURBO DNA-free kit (Invitrogen AM1907). The libraries were prepped using the Illumina Stranded Total RNA Prep, Ligation with Ribo-Zero Plus kit (Illumina 20040529) following the standard protocol. Each library was given a unique index using IDT for Illumina – DNA/RNA UD indexes Set A (Illumina 20026121). The library was pooled in equimolar amounts. Paired-end sequencing on the pooled library was done using the Illumina NextSeq2000 (read length of 100bp and read depth of approximately 24 million reads per library).

### RNA-seq analysis

Sequencing reads (fastq files) were mapped to the reference human genome GRCh38 using STAR aligner version 2.7.10a. The bam files were indexed using samtools version 1.10. RPKM normalized coverage files were generated from the sorted bam files using bamCoverage function of deepTools package version 3.4.5. A read counts table was generated using featureCounts version 1.6.4. The read counts table was used as an input for DESeq2 version 1.40.2 to perform differential gene expression analysis. Transcript isoforms were quantified from fastq files with the quant function from Salmon (version 1.10.3) using the default settings and the human GRCh38 transcriptome. DLD1 parental cells were treated with auxin for 16 and 96 hours to generate a list of auxin sensitive genes. Auxin sensitive genes were manually subtracted from differentially expressed genes from each degron cell line at both 16 and 96 hour time points.

## Supporting information

Merged Supplementary Figures

## Data availability

High-throughput sequencing data generated in this study is available at the NCBI gene expression omnibus (GEO) database under the accession number GSE289252.

