## Supplementary material for "Lamin B1 and LAP2β resist cytoskeletal force to maintain lamin A/C meshwork organization and preserve nuclear integrity": Merged Supplementary Figures

**A.**

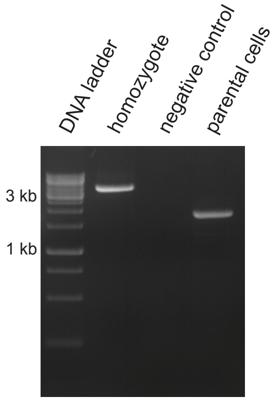


LaminA/C: $F_{out}$ + $R_{out}$
DLD1 cell line

**B.**

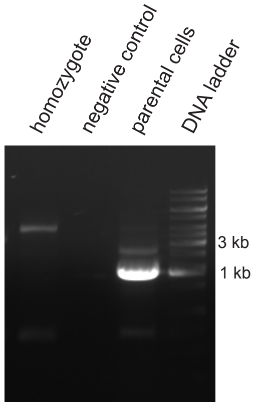


LaminB1: $F_{out2}$ + $R_{out1}$
DLD1 cell line

**C.**

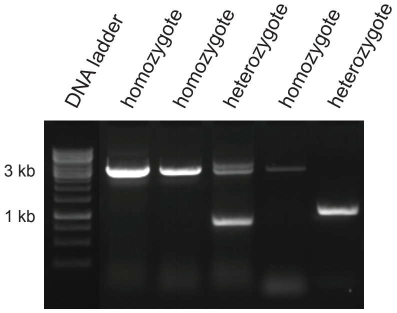


LaminB2: $F_{out1}$ + $R_{out1}$
DLD1 cell line

**Figure S1:** Lamin degron genotyping
(A) Lamin A/C, (B) Lamin B1 and (C) Lamin B2 locus edits were verified by genotyping PCR for homozygous insertion of tags with primers listed in Supplementary Table 4.

**A.**

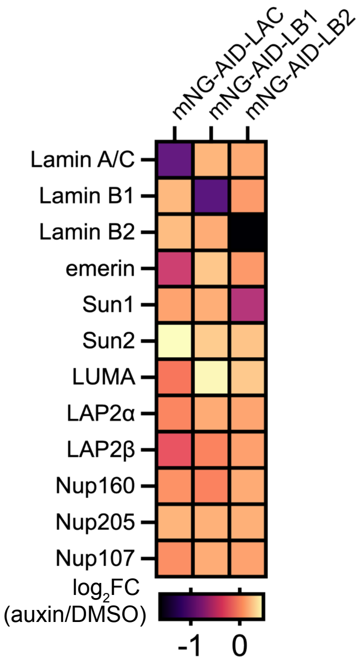
mNG-AID-LAC
mNG-AID-LB1
mNG-AID-LB2
Lamin A/C
Lamin B1
Lamin B2
emerin
Sun1
Sun2
LUMA
LAP2α
LAP2β
Nup160
Nup205
Nup107
log2FC
(auxin/DMSO)
-1
0

**Figure S2:** Additional lamin degron nuclear extract proteomics

(A) Quantification of nuclear envelope protein abundance by multiplexed tandem mass tagging (TMT) in nuclear extracts from mNG-AID-LA/C, mNG-AID-LB1 and mNG-AID-LB2 cells treated with auxin for 12 hours.

**A.** Principal component analysis

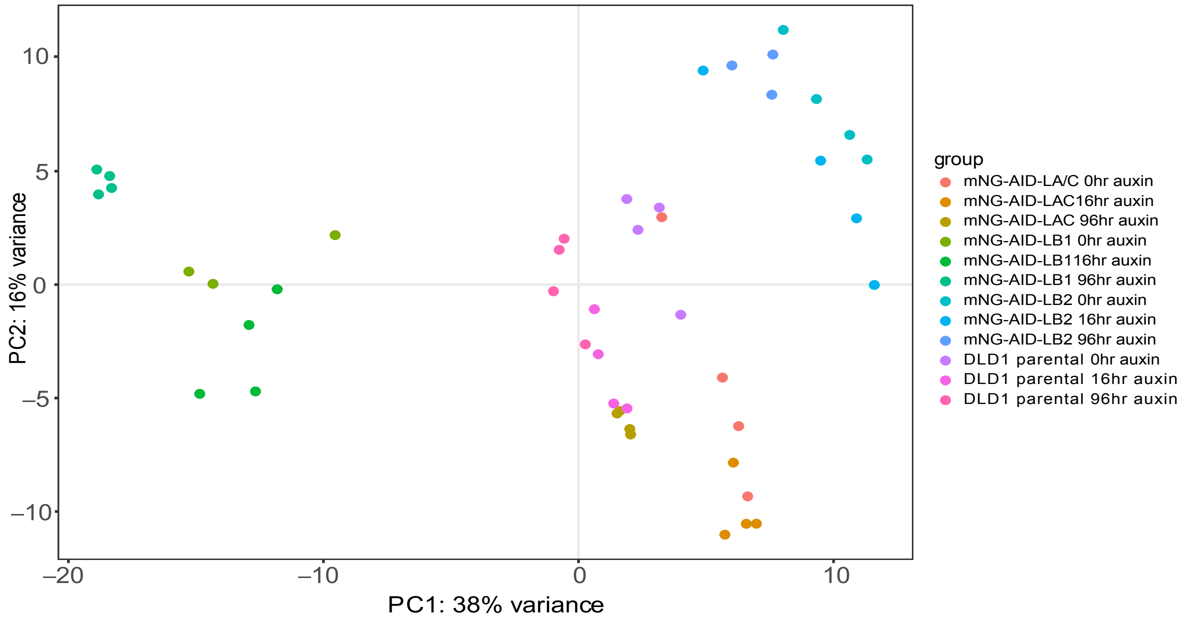


**B.** venn diagram of overlap of differentially expressed genes

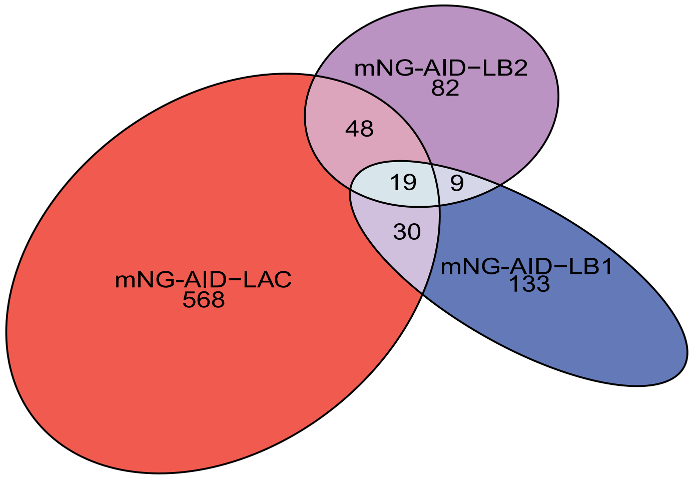


**C.** lamin abundance in DLD1 cells compared to mNG-AID-LA/C, mNG-AID-LB1 or mNG-AID-LB2 cells

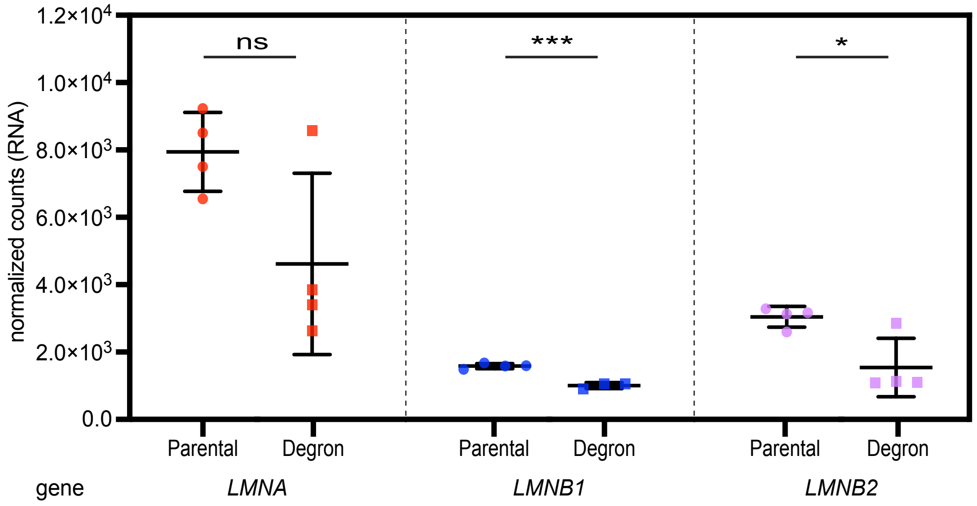

**Figure S3:** RNAseq additional analysis

(A) Principal component analysis of RNAseq data. (B) Venn diagram representing the number of differentially expressed genes that overlapped between mNG-AID-LA/C, mNG-AID-LB1 and mNG-AID-LB2 cells after 96 hours of auxin treatment. (C) *LMNA, LMNB1* or *LMNB2* levels (normalized counts) in DLD1 parental cells compared to AID-tagged cells. * indicates $p_{adj} < 0.05$, *** indicates $p_{adj} < 0.001$ and ns indicates $p_{adj} > 0.05$.

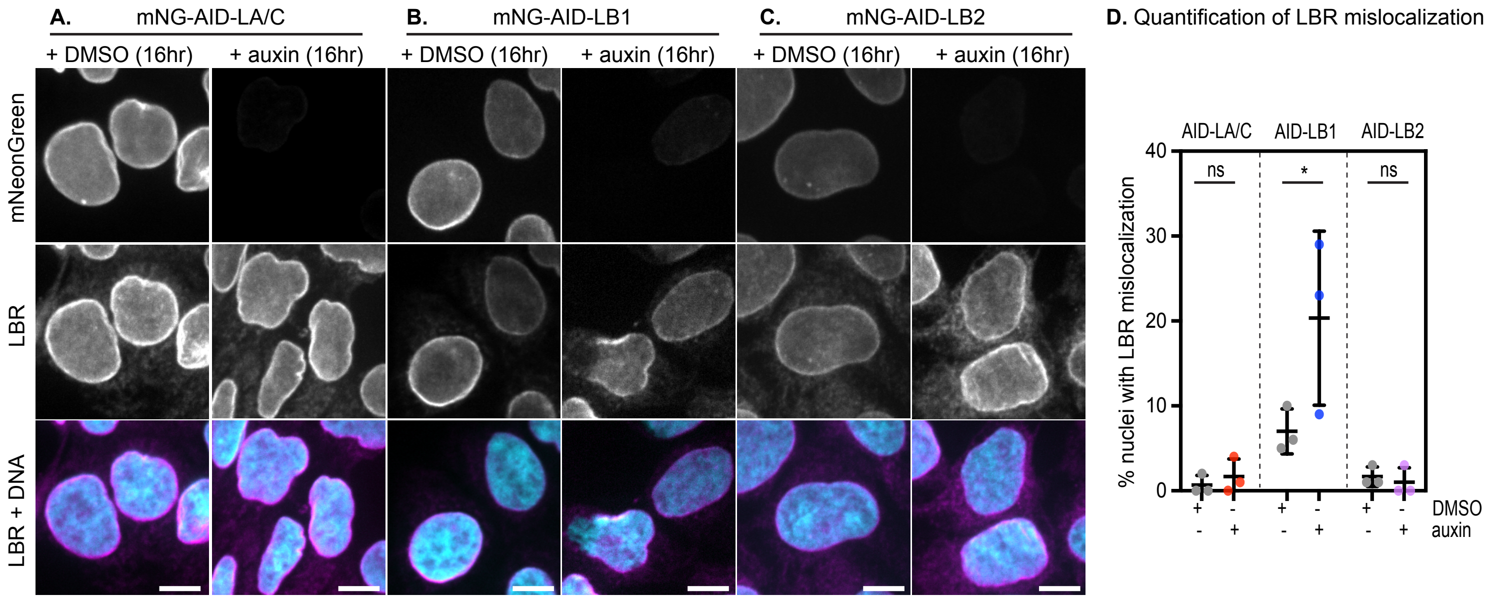

A.
mNG-AID-LA/C
B.
mNG-AID-LB1
C.
mNG-AID-LB2
+ DMSO (16hr)
+ auxin (16hr)
mNeonGreen
LBR
LBR + DNA
D. Quantification of LBR mislocalization
AID-LA/C
AID-LB1
AID-LB2
ns
*
% nuclei with LBR mislocalization
0
10
20
30
40
DMSO
auxin

**Figure S4:** Lamin B1 depletion results in mislocalization of LBR
(A-C) Immunofluorescence of mNeonGreen and LBR after 16 hours of DMSO or auxin treatment in mNG-AID-LA/C (A), mNG-AID-LB1 (B), and mNG-AID-LB2 (C) cells. Scale bar, 10 μm. (D) Percentage of nuclei with LBR mislocalization after 16 hours of auxin treatment in AID-LAC, AID-LB1 and AID-LB2 cells. * indicates $p < 0.05$ and ns indicates $p > 0.05$ by pairwise t-test. Average values of 3 independent replicate experiments shown; $n > 400$ cells analyzed per condition.

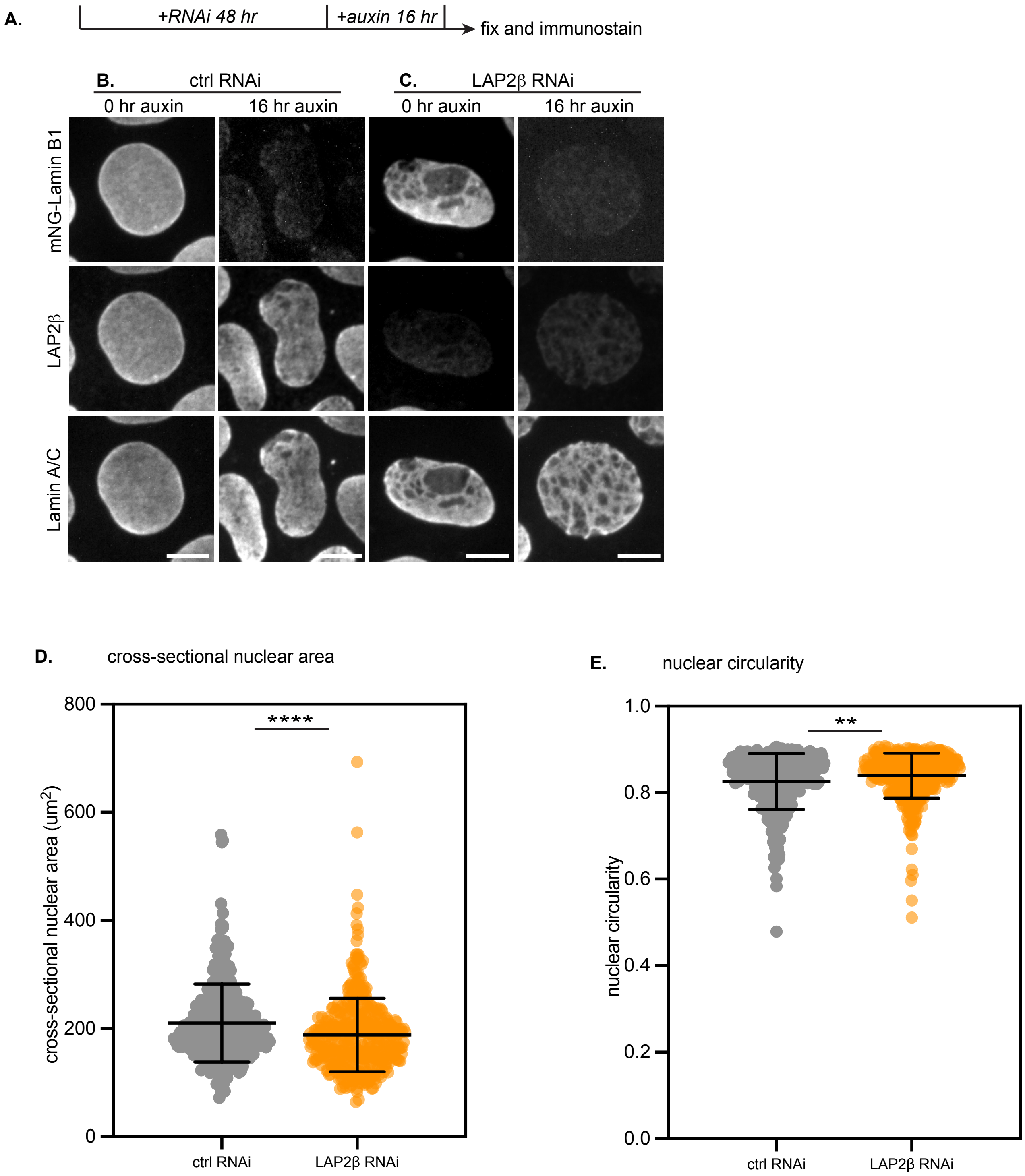

A.
+RNAi 48 hr
+auxin 16 hr
fix and immunostain
B.
ctrl RNAi
C.
LAP2β RNAi
0 hr auxin
16 hr auxin
0 hr auxin
16 hr auxin
mNG-Lamin B1
LAP2β
Lamin A/C
D.
cross-sectional nuclear area
cross-sectional nuclear area (um²)
0
200
400
600
800
****
ctrl RNAi
LAP2β RNAi
E.
nuclear circularity
nuclear circularity
0.0
0.2
0.4
0.6
0.8
1.0
**
ctrl RNAi
LAP2β RNAi

**Figure S5:** LAP2β depletion in mNG-AID-LB1 cells is sufficient to drive changes in nuclear morphology.

(A) Overview of experimental strategy. mNG-AID-LB1 cells were treated with control or LAP2β RNAi followed by DMSO or auxin for 16 hours. mNG-AID-LB1 cells were then fixed and immunostained. (B-C) Immunofluorescence of mNeonGreen, LAP2β , and lamin A/C after mNG-AID-LB1 cells were treated with (B) non-targeting control or (C) LAP2β RNAi for 48 hours. Scale bar, 5 μm (D) Cross-sectional nuclear area and (E) nuclear circularity after mNG-AID-LB1 cells were subject to non-targeting control or LAP2β RNAi. $n > 300$ cells analyzed per condition from 3 independent replicate experiments. ** indicates $p < 0.01$ and **** indicates $p < 0.0001$ by pairwise t-test.

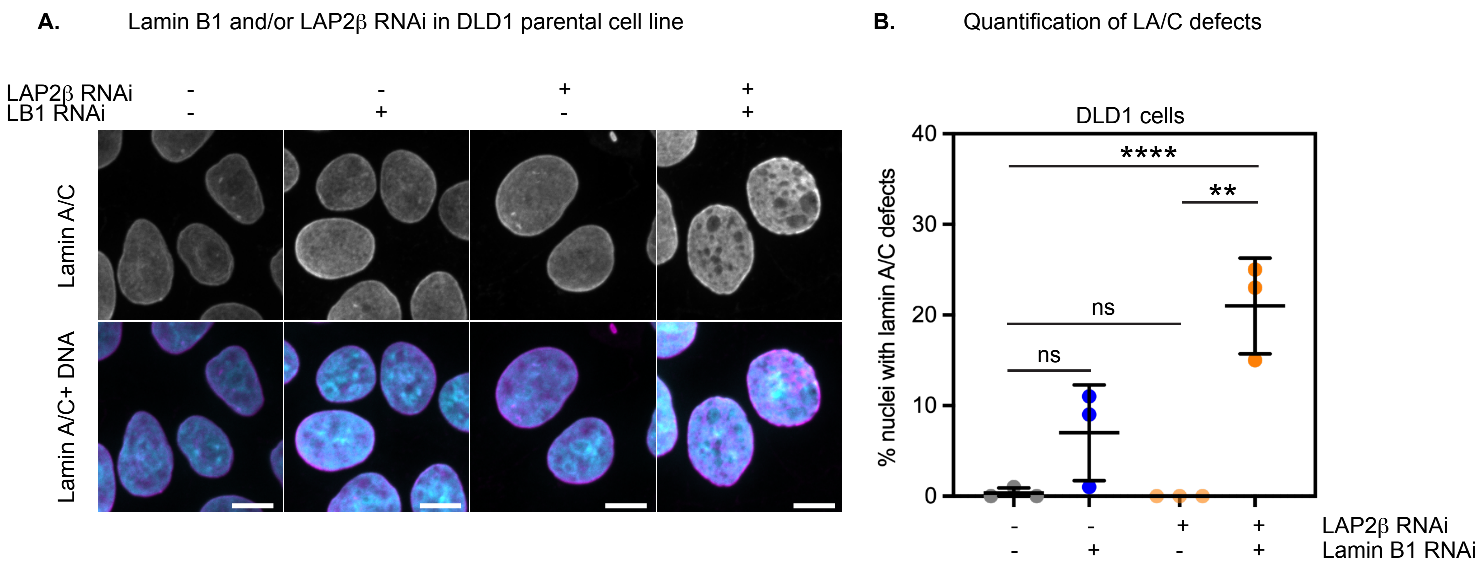
A.
Lamin B1 and/or LAP2β RNAi in DLD1 parental cell line
LAP2β RNAi
LB1 RNAi
Lamin A/C
Lamin A/C+ DNA
B.
Quantification of LA/C defects
DLD1 cells
% nuclei with lamin A/C defects
40
30
20
10
0
****
**
ns
ns
LAP2β RNAi
Lamin B1 RNAi

**Figure S6:** LAP2β and Lamin B1 co-depletion increases lamin A/C meshwork defects in DLD1 parental cell line

(A) Immunofluorescence of lamin A/C in DLD1 parental cells after control RNAi, lamin B1 RNAi, LAP2β RNAi, or lamin B1 and LAP2β RNAi. Scale bar, 10 μm. (B) Quantification of the percentage of nuclei with lamin A/C defects in conditions shown in (A). Each dot on graph represents one experimental replicate; $n > 350$ cells analyzed per condition across 3 replicate experiments. ns indicates $p > 0.05$, ** indicates $p < 0.01$ and **** indicates $p < 0.0001$ by one way ANOVA.
